# Cryo-EM structure of a eukaryotic zinc transporter at a low pH suggests its Zn^2+^-releasing mechanism

**DOI:** 10.1101/2022.06.29.497979

**Authors:** Senfeng Zhang, Chunting Fu, Yongbo Luo, Qingrong Xie, Tong Xu, Ziyi Sun, Zhaoming Su, Xiaoming Zhou

**Affiliations:** Department of Integrated Traditional Chinese and Western Medicine, Rare Diseases Center, State Key Laboratory of Biotherapy, West China Hospital, Sichuan University, Chengdu, Sichuan 610041, China; The State Key Laboratory of Biotherapy, Department of Geriatrics and National Clinical Research Center for Geriatrics, West China Hospital, Sichuan University, Chengdu, Sichuan 610041, China

**Keywords:** ZnT8, SLC30, CDF, Protonation, Cryo-EM structure, Single-particle reconstruction, Membrane protein, Transport cycle, Conformational change

## Abstract

Zinc transporter 8 (ZnT8) is mainly expressed in pancreatic islet β cells and is responsible for H^+^-coupled uptake (antiport) of Zn^2+^ into the lumen of insulin secretory granules. Structures of human ZnT8 and its prokaryotic homolog YiiP have provided structural basis for constructing a plausible transport cycle for Zn^2+^. However, the mechanistic role that protons play in the transport process remains unclear. Here we present a lumen-facing cryo-EM structure of ZnT8 from *Xenopus tropicalis* (xtZnT8) in the presence of Zn^2+^ at a luminal pH (5.5). Compared to a Zn^2+^-bound xtZnT8 structure at a cytosolic pH (7.5), the low-pH structure displays an empty transmembrane Zn^2+^-binding site with a disrupted coordination geometry. Combined with a Zn^2+^-binding assay our data suggest that protons may disrupt Zn^2+^ coordination at the transmembrane Zn^2+^-binding site in the lumen-facing state, thus facilitating Zn^2+^ release from ZnT8 into the lumen.

## Introduction

Zinc is the second most abundant trace metal in cells and plays critical roles in many biological processes, including cell growth and development, functioning of the central nervous system and the immune system (Kambe et al., 2015; Maret, 2013; Vallee and Falchuk, 1993). A variety of enzymes also require Zn^2+^ for their biological functions (Kambe et al., 2015; Maret, 2013; Vallee and Falchuk, 1993). Meanwhile, excessive cytoplasmic free Zn^2+^ is highly toxic. Therefore, the intracellular homeostasis of Zn^2+^ is tightly controlled, mainly by two classes of solute carrier (SLC) transporters: the ZRT/IRT-like proteins (ZIPs, or SLC39), and the cation-diffusion facilitators (CDFs, or SLC30), also known as zinc transporters (ZnTs) (Bin et al., 2018; Eide, 2006; Kambe et al., 2015). These two transporter families mediate influx and efflux of Zn^2+^ into and out of the cytoplasm of the cell, respectively. Among ZnTs, ZnT8 is mainly expressed in pancreatic islet β cells and is responsible for H^+^-coupled uptake/antiport of Zn^2+^ into the lumen of insulin secretory granules, in which Zn^2+^ is complexed with insulin in a crystalline form and is co-secreted with insulin (Davidson et al., 2014). ZnT8 is often targeted by autoantibodies in type 1 diabetes (Kawasaki, 2012; Wenzlau et al., 2007), whereas mutations and single nucleotide polymorphisms of ZnT8 are associated with type 2 diabetes (Flannick et al., 2014; Saxena et al., 2007; Scott et al., 2007; Sladek et al., 2007; Zeggini et al., 2007).

High-resolution structures of the ZnT family were first obtained with its prokaryotic homolog, YiiP from *Escherichia coli* (ecYiiP) (Lu and Fu, 2007; Lu et al., 2009) or *Shewanella oneidensis* (soYiiP) (Coudray et al., 2013; Lopez-Redondo et al., 2021; Lopez-Redondo et al., 2018), revealing a homodimeric configuration of YiiP. Recently, structures of human ZnT8 (hZnT8) were determined by single-particle cryo-electron microscopy (cryo-EM) (Xue et al., 2020), also as a homodimer (Fig. 1A). Four Zn^2+^-binding sites were identified in each hZnT8 protomer (Xue et al., 2020): one S_TMD_ site at the center of the transmembrane domain (TMD), two adjacent S_CTD_ sites in the carboxy-terminal domain (CTD), and one S_interface_ site at the interface between TMD and CTD (Fig. 1A). S_TMD_ is the primary site for Zn^2+^ binding and transport, whereas the S_CTD_ sites are formed by residues from both protomers and contribute to hZnT8 dimerization (Xue et al., 2020). These three Zn^2+^-binding sites are highly conserved in the ZnT family (Fig. 2), including the prokaryotic YiiP (Lopez-Redondo et al., 2018; Lu and Fu, 2007). On the other hand, the S_interface_ site seems less conserved. In hZnT8, it is formed by residues from both CTD and a loop region between TM2 and TM3 (Xue et al., 2020), while in YiiP it is formed solely by residues from the TM2-TM3 loop (Lopez-Redondo et al., 2018; Lu and Fu, 2007). In some species like mice and rats, no Zn^2+^-coordinating residue (Cys, His, Glu or Asp) is present within the predicted TM2-TM3 loop (Fig. 2), further suggesting that S_interface_ may not be conserved. Nonetheless, it has been hypothesized that S_interface_ may increase the local concentration of Zn^2+^to facilitate its binding to the S_TMD_ site in ZnT8 (Xue et al., 2020).

**Figure 1.**
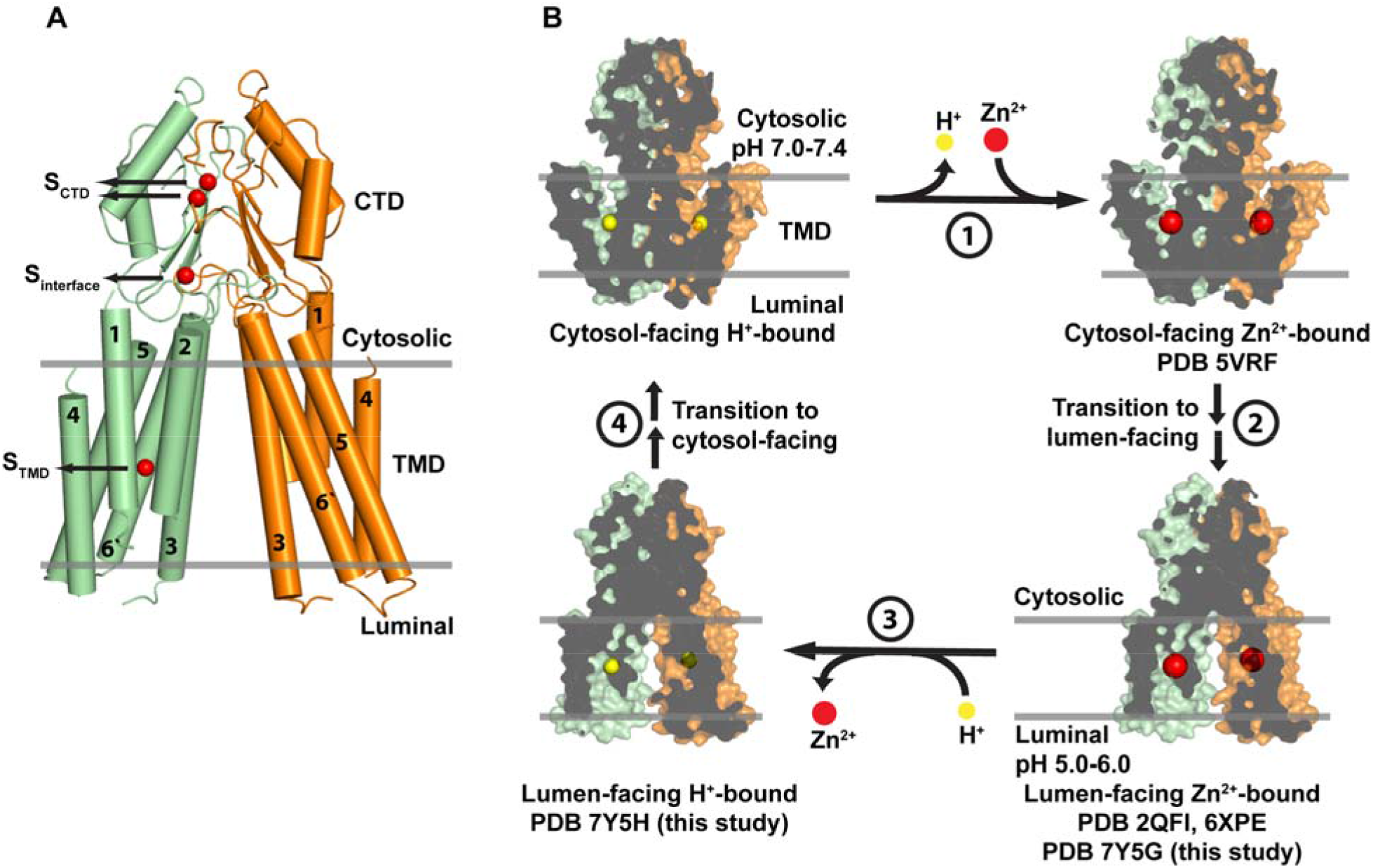
Structure and a working model for Zn^2+^ transport in ZnT8. (A) Cryo-EM structure of hZnT8 in the presence of Zn^2+^ in a lumen-facing conformation (PDB: 6XPE). The hZnT8 dimer is displayed in cartoon mode with two protomers colored in green and orange, respectively. Bound Zn^2+^ are rendered as red spheres in one protomer only for better viewing. Transmembrane segments of each protomer are numbered from one to six. The relative position of the granule membrane is indicated by two grey lines. (B) A simplified alternating-access model for ZnT8. The dimeric ZnT8 models are displayed in surface mode with two protomers colored in green and orange, respectively. S_TMD_-bound H^+^/Zn^2+^ are rendered as spheres as indicated.

**Figure 2.**
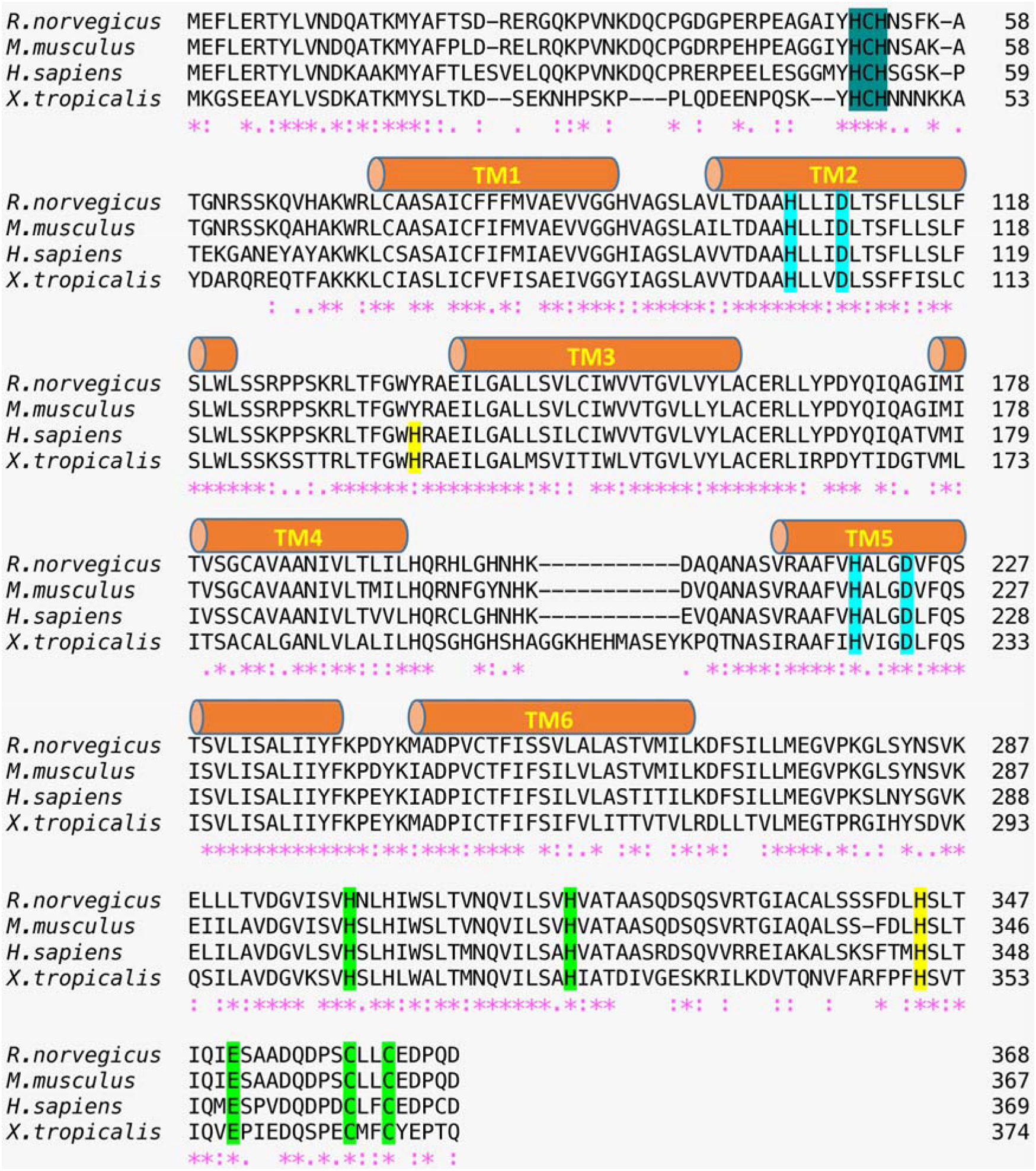
Sequence alignment of ZnT8 homologs by ClustalW (Combet et al., 2000; Thompson et al., 1994). Transmembrane segments are indicated by cylinders and labeled from TM1 to TM6. The S_TMD_ site residues are highlighted in cyan, the S_CTD_ sites in green (including the HCH motif in dark green), and the S_interface_ site in yellow. Asterisks (*) indicate identical residues. Colons (:) indicate strong similarities. Periods (.) indicate weak similarities. *R. norvegicus, Rattus norvegicus; M. musculus, Mus musculus; H. sapiens, Homo sapiens; X. tropicalis, Xenopus tropicalis*.

The structures of hZnT8 and YiiP were captured in both outward-facing (lumen-facing) and inward-facing (cytosol-facing) states (Lopez-Redondo et al., 2018; Lu and Fu, 2007; Xue et al., 2020), allowing construction of an alternating-access model for ZnT8 transporters as follows (Fig. 1B). (1) Due to a slightly basic cytosolic pH (Llopis et al., 1998; Stiernet et al., 2007), a counter-transported H^+^ is released to the cytosol from ZnT8 in the cytosol-facing state, allowing a cytosolic Zn^2+^ to bind to the S_TMD_ site. (2) ZnT8 transits to the lumen-facing state. (3) The S_TMD_-bound Zn^2+^ is released to the granule lumen while the acidic luminal pH (Hutton, 1982) protonates the S_TMD_ site of ZnT8. (4) ZnT8 transits back to the cytosol-facing state, ready for the next cycle. The model is consistent with the structural and functional data reported for ZnTs regarding H^+^-coupled Zn^2+^ binding and transport (Ohana et al., 2009; Shusterman et al., 2014; Xue et al., 2020). However, the role of H^+^ binding in this process (i.e. step 3) remains speculative.

In this study, we compared two lumen-facing cryo-EM structures of ZnT8 from *Xenopus tropicalis* (xtZnT8) in the presence of Zn^2+^ at either a cytosolic pH (7.5) or a luminal pH (5.5) to assess the effect of H^+^ binding on ZnT8 in the lumen-facing state. Together with a Zn^2+^-binding analysis our data suggest a role of protons that may facilitate the release of Zn^2+^ from ZnT8 into the lumen.

## Results

### Cryo-EM structure of xtZnT8 with Zn^2+^ at a cytosolic pH (7.5) in a lumen-facing state

First, we determined the structure of xtZnT8 in the presence of 1 mM Zn^2+^ at a cytosolic pH of 7.5 (3.9 Å, xtZnT8-Zn^2+^) by cryo-EM (Fig. 3). XtZnT8 shares 58% sequence identity and 83% sequence similarity to hZnT8 (Fig. 2). Similar to the Zn^2+^-bound hZnT8 structure (hZnT8-Zn^2+^, PDB: 6XPE), the xtZnT8-Zn^2+^ structure is also organized as a homodimer (Fig. 4A). Structural alignment between individual protomers of xtZnT8-Zn^2+^ and hZnT8-Zn^2+^yielded a root mean square deviation (RMSD) of 1.18 Å with all C_α_ atoms aligned, while the major deviation came from flexible loop regions of the two proteins, indicating that the two structures are highly alike (Fig. 4A). Similarly, each protomer of xtZnT8-Zn^2+^ has two adjacent S_CTD_ sites (S_CTD1_ and S_CTD2_), which are formed by the HCH (His45-Cys46-His47) motif from one protomer and multiple Zn^2+^-coordinating residues from the other protomer, occupied by two Zn^2+^ (Fig. 2 and 4B). Also, the xtZnT8-Zn^2+^ structure has one Zn^2+^ bound at the S_TMD_ site with a tetrahedral coordination, which is formed by His100 and Asp104 from TM2, and His225 and Asp229 from TM5 (Fig. 2 and 4C). Consistently, the S_TMD_ residues are highly conserved in eukaryotic species with the 2-His-2-Asp configuration (Cotrim et al., 2019; Hoch et al., 2012) (Fig. 2). Like the hZnT8-Zn^2+^ structure, the S_TMD_-bound Zn^2+^ in xtZnT8-Zn^2+^ is accessible by solvent from the luminal side (Fig. 4D), indicating that the transporter is lumen-facing. Therefore, the xtZnT8-Zn^2+^ structure was also captured in a Zn^2+^-bound lumen-facing state similar to hZnT8-Zn^2+^ (Xue et al., 2020). Interestingly, the S_TM_D-bound Zn^2+^ was not released automatically from the S_TMD_ site in the lumen-facing state, suggesting that a Zn^2+^-releasing mechanism is required for eukaryotic ZnT8.

**Figure 3.**
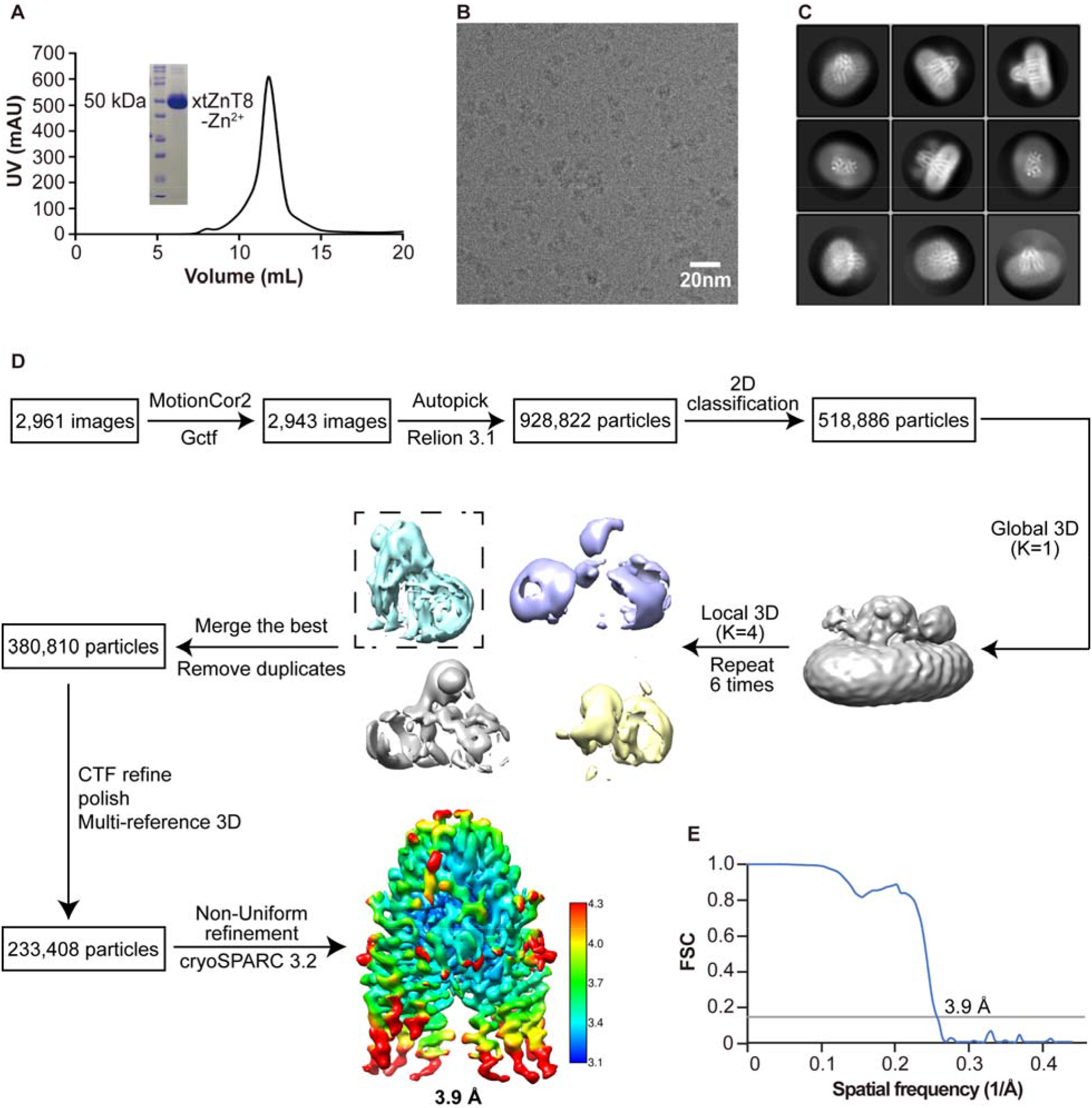
Cryo-EM sample preparation and data processing of xtZnT8-Zn^2+^. (A) The elution profile of xtZnT8-Zn^2+^ in the presence of 1 mM Zn^2+^at pH 7.5 on a size-exclusion column. The insert shows SDS-PAGE analysis of the purified sample. (B) A representative area of cryo-EM micrographs of xtZnT8-Zn^2+^ particles. (C) Selected 2D class averages of xtZnT8-Zn^2+^ particles. (D) The workflow of cryo-EM data processing of the xtZnT8-Zn^2+^ dataset. (E) The overall resolution of xtZnT8-Zn^2+^ was determined by the “gold standard” FSC curve using the FSC=0.143 criterion.

**Figure 4.**
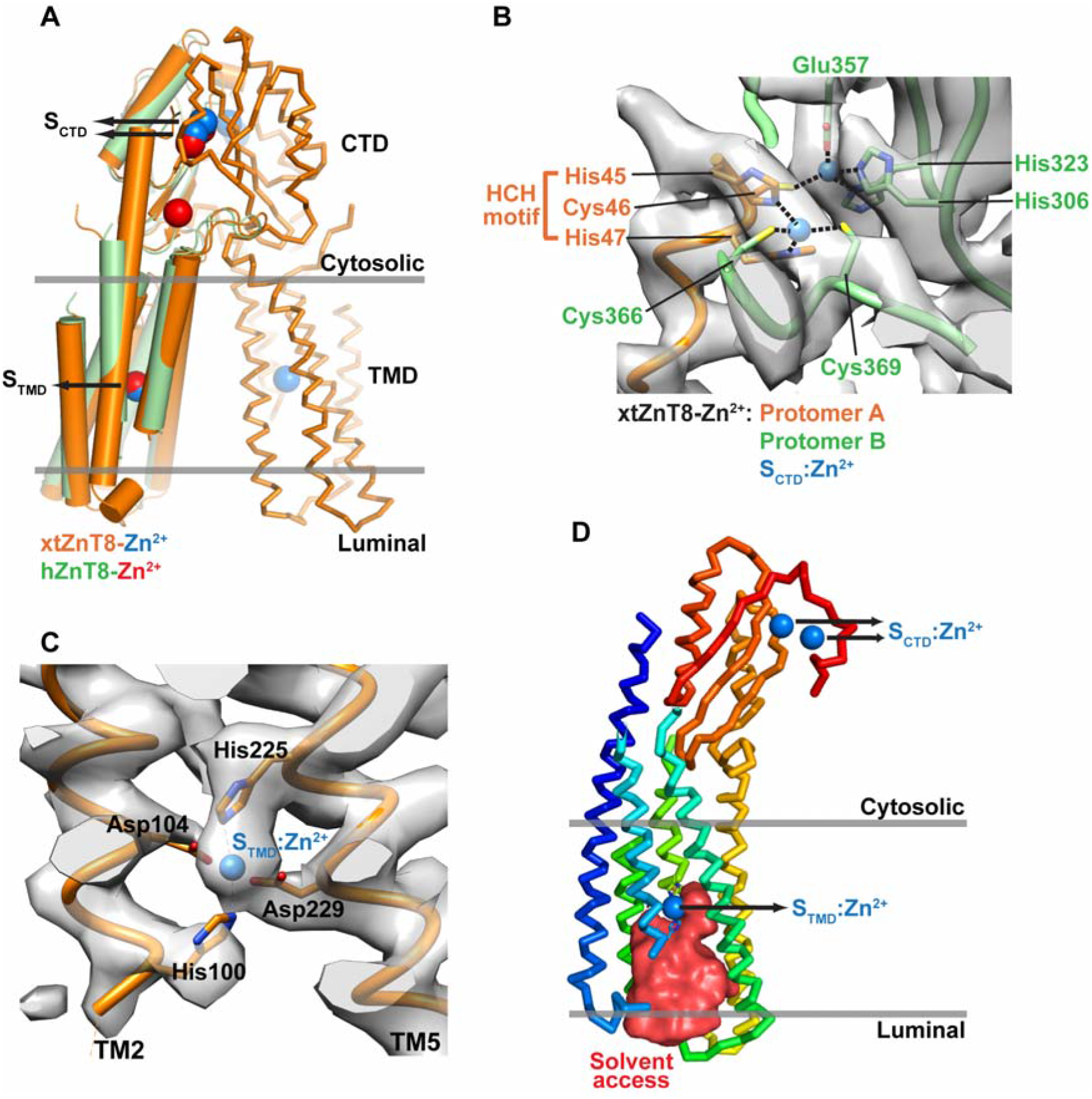
Cryo-EM structure of xtZnT8-Zn^2+^ in a lumen-facing state. (A) The xtZnT8-Zn^2+^ dimer is displayed in orange, with one protomer in cartoon mode and the other as ribbons. Bound Zn^2+^ are shown as blue spheres. One protomer of the hZnT8-Zn^2+^ structure (PDB: 6XPE) is displayed in green with bound Zn^2+^ rendered as red spheres, and is superimposed onto xtZnT8-Zn^2+^. The relative position of the granule membrane is indicated by two grey lines. (B) The S_CTD_ sites of xtZnT8-Zn^2+^. Two protomers are displayed as ribbons with one in orange and the other in green. The S_CTD_-bound Zn^2+^ are shown as blue spheres. The Zn^2+^-coordinating residues are displayed as sticks, and the HCH motif is indicated. The cryo-EM map of xtZnT8-Zn^2+^ is shown as grey densities. (C) One Zn^2+^ (blue sphere) binds to the S_TMD_ site in xtZnT8-Zn^2+^. The Zn^2+^-coordinating residues are displayed as sticks. The cryo-EM map of xtZnT8-Zn^2+^ is shown as grey densities. (D) One protomer of xtZnT8-Zn^2+^ is displayed as ribbons, which is colored in rainbow spectrum with blue for the amino-terminus. Bound Zn^2+^ are shown as blue spheres. The solvent-accessible space is shown as a red surface. The relative position of the granule membrane is indicated by two grey lines.

### A lumen-facing cryo-EM structure of xtZnT8 at a luminal pH (5.5)

It has been assumed that binding of luminal protons to the S_TMD_ site (likely histidine residues) in a Zn^2+^-bound lumen-facing ZnT8 would weaken Zn^2+^ binding and promote Zn^2+^ release to the lumen (Xue et al., 2020) due to an acidic pH (5~6) of insulin secretory granules (Hutton, 1982). Similar assumptions have been proposed for YiiPs as a Zn^2+^-releasing mechanism (Gupta et al., 2014; Lopez-Redondo et al., 2018). However, structural evidence to support this hypothesis is lacking. To investigate the effect of protons on Zn^2+^ binding in ZnT8, we then determined another cryo-EM structure of xtZnT8 in the presence of 1 mM Zn^2+^ at a luminal pH of 5.5 (3.7 Å, xtZnT8-H^+^) (Fig. 5). The xtZnT8-H^+^ sample was prepared in the same way as the xtZnT8-Zn^2+^ sample except that the former underwent an additional buffer exchange step from pH 7.5 to pH 5.5. Ideally, this experimental design would allow us to assess the effect of protons when they bind to xtZnT8-Zn^2+^.

**Figure 5.**
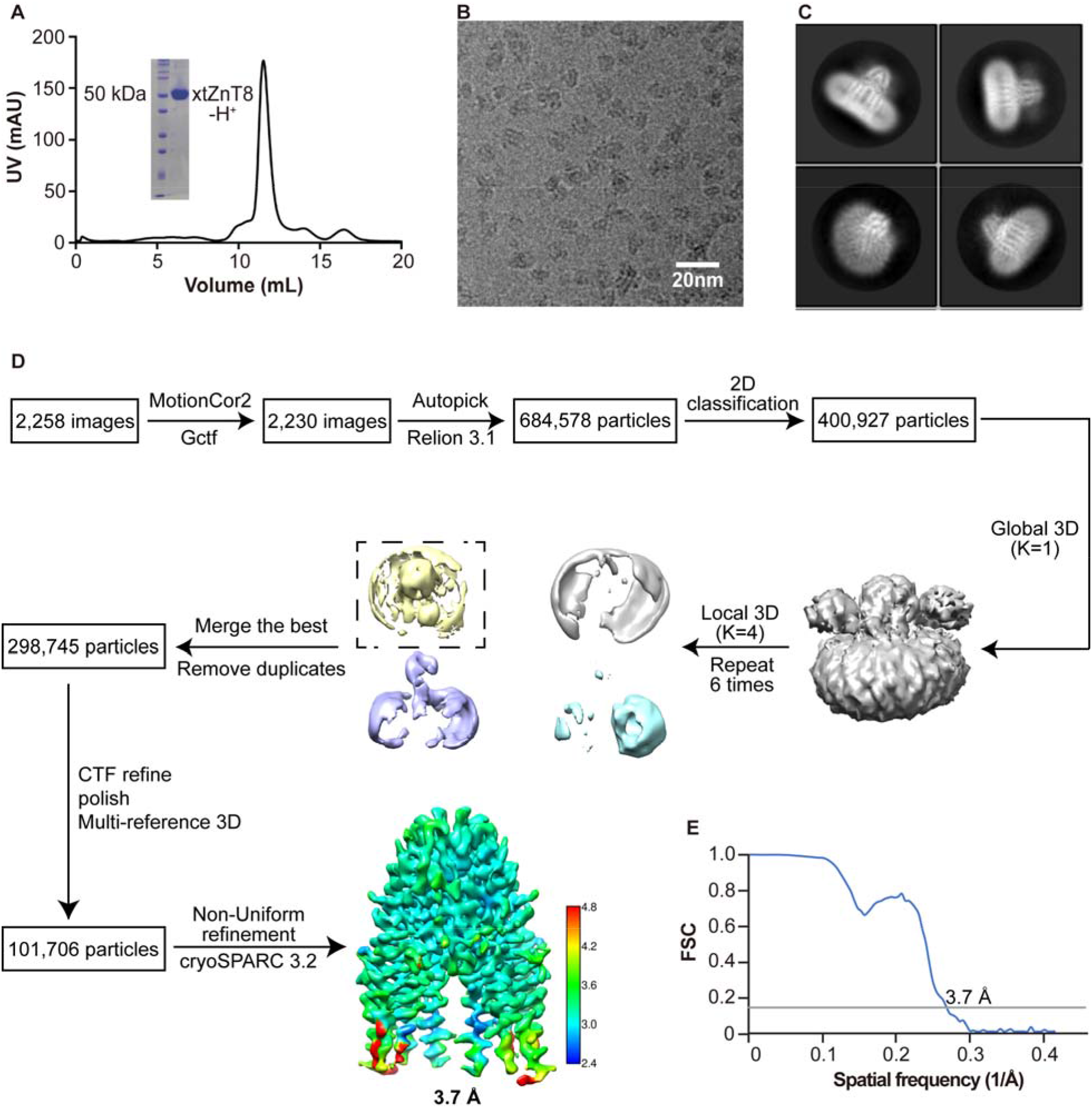
Cryo-EM sample preparation and data processing of xtZnT8-H. (A) The elution profile of xtZnT8-H^+^ in the presence of 1 mM Zn^2+^ at pH 5.5 on a size-exclusion column. The insert shows SDS-PAGE analysis of the purified sample. (B) A representative area of cryo-EM micrographs of xtZnT8-H^+^ particles. (C) Selected 2D class averages of xtZnT8-H^+^ particles. (D) The workflow of cryo-EM data processing of the xtZnT8-H^+^ dataset. (E) The overall resolution of xtZnT8-H^+^ was determined by the “gold standard” FSC curve using the FSC=0.143 criterion.

Interestingly, the xtZnT8-H^+^ structure also adopts a lumen-facing conformation, similar to xtZnT8-Zn^2+^ (Fig. 6A). Superposition of the two xtZnT8 structures yielded an all-atom RMSD of 0.58 Å, indicating that xtZnT8-H^+^ and xtZnT8-Zn^2+^ are very alike (Fig. 6A). However, one prominent difference between the two structures is that the side chain of His100 in xtZnT8-H^+^ is pointing toward the luminal side, thus disrupting the tetrahedral coordination for Zn^2+^ at the S_TMD_ site (Fig. 6B). Consistently, no Zn^2+^ density was observed at S_TMD_ in xtZnT8-H^+^ even in the presence of 1 mM Zn^2+^ (Fig. 6C). This data suggests that exposure of xtZnT8-Zn^2+^ to protons at a luminal pH causes release of the bound Zn^2+^ from the S_TMD_ site, providing structural evidence to support protonation as a Zn^2+^-releasing mechanism in ZnT8.

**Figure 6.**
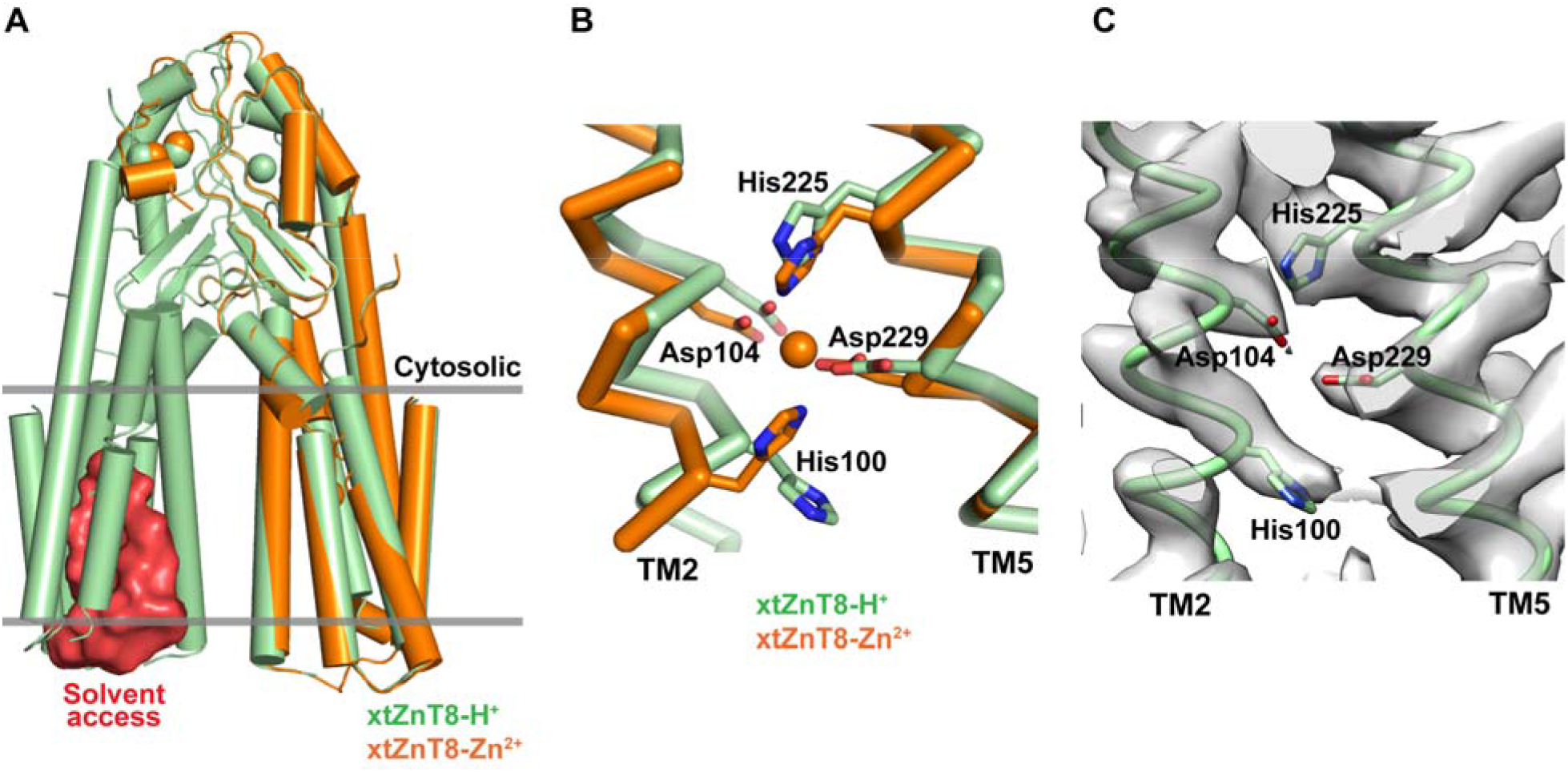
Cryo-EM structure of xtZnT8-H* in a lumen-facing state. (A) The xtZnT8-H^+^ dimer is displayed in green, with a red surface indicating the solvent-accessible space for one protomer. One protomer of the xtZnT8-Zn^2+^ structure is displayed in orange and is superimposed onto xtZnT8-H^+^. Bound Zn^2+^ are shown as spheres. The relative position of the granule membrane is indicated by two grey lines. (B) Comparison of the S_TMD_ site between xtZnT8-H^+^ (in green) and xtZnT8-Zn^2+^ (in orange). One Zn^2+^ (orange sphere) binds to S_TMD_ in xtZnT8-Zn^2+^. The Zn^2+^-coordinating residues are rendered as sticks. (C) The cryo-EM density map of the S_TMD_ site in xtZnT8-H^+^. The Zn^2+^-coordinating residues are rendered as sticks.

### Protons disrupt Zn^2+^ binding in xtZnT8

A functional inference from the xtZnT8-H^+^ structure is that protons at a luminal pH (e.g. 5.5) would disrupt Zn^2+^ binding to xtZnT8. To test this idea, we used microscale thermophoresis (MST) (Jerabek-Willemsen et al., 2011) to measure Zn^2+^ binding to xtZnT8 in various conditions. First, we titrated Zn^2+^ to wild-type xtZnT8 purified in the absence of Zn^2+^ at a cytosolic pH of 7.5. The MST data was readily fit by an inverse S-shaped curve (Fig. 7A), with an equilibrium dissociation constant *(Kd)* of 4.91 ± 0.39 μM for Zn^2+^. The titrated Zn^2+^ binding can be attributed to the S_TMD_ site of xtZnT8, as an S_T_MD-double-mutant (H100A/H225A) showed a noise-like trendless no-binding pattern when Zn^2+^ was titrated (Fig. 7B). We then titrated Zn^2+^ to wild-type xtZnT8 purified in the absence of Zn^2+^ at a luminal pH of 5.5 and obtained a noise-like no-binding pattern as well (Fig. 7C), suggesting that protons at a low pH disrupt Zn^2+^ binding to the S_TMD_ site of xtZnT8. These data are consistent with the xtZnT8 structures.

**Figure 7.**
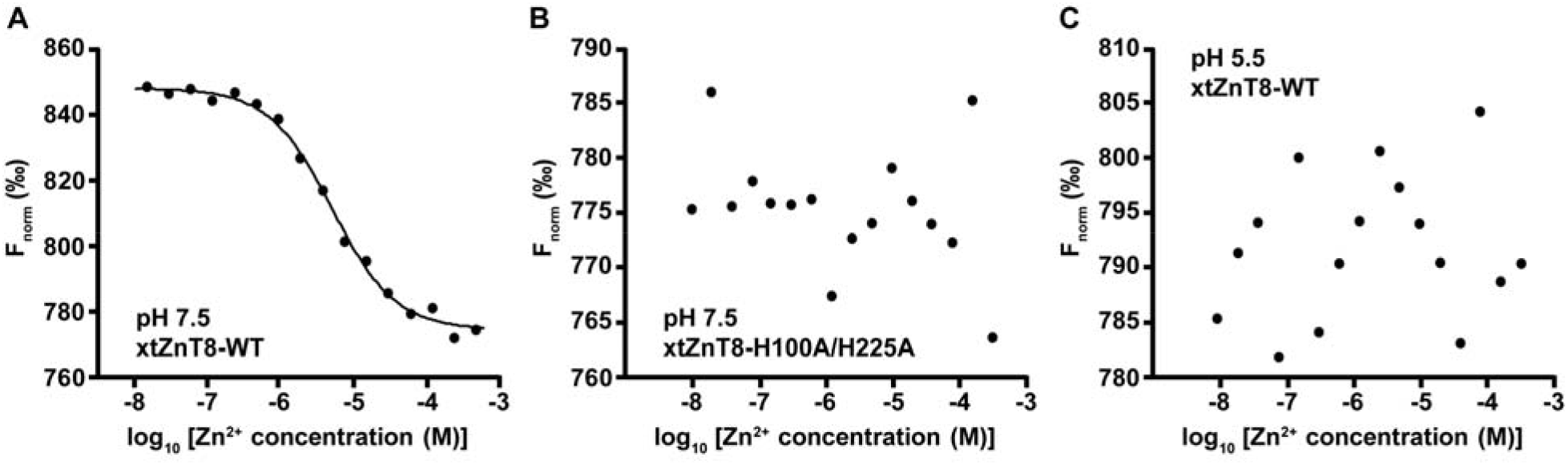
Binding of Zn^2+^ to xtZnT8 measured by microscale thermophoresis. (A) A representative MST curve for wild-type xtZnT8 (xtZnT8-WT) titrated with Zn^2+^ at pH 7.5. (B) A representative MST measurement for an S_TMD_-double-mutant of xtZnT8 (xtZnT8-H100A/H225A) titrated with Zn^2+^ at pH 7.5. (C) A representative MST measurement for wild-type xtZnT8 titrated with Zn^2+^ at pH 5.5. The detailed procedure of MST measurements were described in the “Materials and methods” section.

## Discussion

### A potential Zn^2+^-releasing mechanism in ZnT8

Another major difference between the two xtZnT8 structures is that a portion of TM2 (Ala93-Ala98), which is close to the luminal side, appears more flexible in xtZnT8-Zn^2+^ than in xtZnT8-H^+^, and therefore is less visible in the former map (Fig. 8A). This is reminiscent of hZnT8-Zn^2+^ (PDB: 6XPE) and the structure of its S_T_MD-double-mutant (hZnT8-D110N/D224N, PDB: 6XPD), which mimics a protonated state at positions 110 and 224 and doesn’t bind Zn^2+^ at the mutated S_TMD_ site (Xue et al., 2020). Compared to hZnT8-D110N/D224N, the hZnT8-Zn^2+^ structure also shows a less defined TM2 portion near the luminal side (Fig. 8B). This result suggests that in the lumen-facing state of ZnT8, binding of Zn^2+^ at S_TMD_ may increase the flexibility of the near-lumen TM2 region, thus allowing or increasing solvent access from the luminal side. Therefore, a potential Zn^2+^-releasing mechanism for ZnT8 can be proposed as in Fig. 8C. Briefly, in the Zn^2+^-bound lumen-facing state of xtZnT8, the TM2 region near the luminal side becomes flexible, allowing luminal protons (pH 5~6) to access and protonate the S_TMD_ site (likely His100/His225), which disrupts Zn^2+^ coordination and facilitates Zn^2+^ release from S_TMD_ to the lumen.

**Figure 8.**
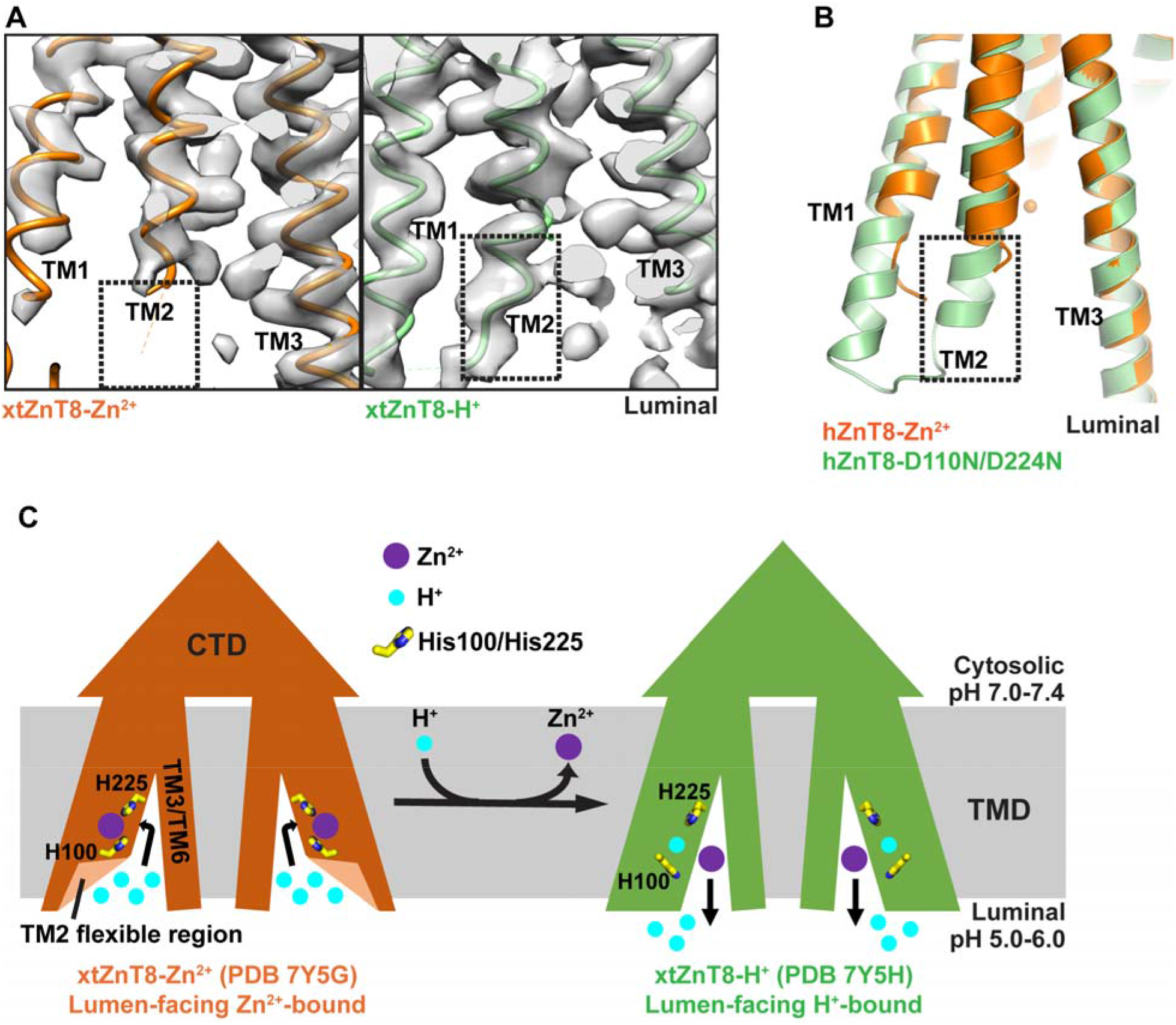
A potential Zn^2+^-releasing mechanism in ZnT8. (A) Comparison of the cryo-EM density maps between xtZnT8-Zn^2+^ (orange ribbons, left) and xtZnT8-H^+^ (green ribbons, right) for a TM2 region near the luminal side (boxed). (B) Structural comparison between hZnT8-Zn^2+^ (orange ribbons) and hZnT8-DHON/D224N (green ribbons) for a TM2 region near the luminal side (boxed). (C) A l-Γ-facilitated Zn^2+^-releasing mechanism is proposed for ZnT8 based on xtZnT8-Zn^2+^ and xtZnT8-H^+^ structures. Briefly, in the Zn^2+^-bound lumen-facing state of ZnT8, a flexible TM2 region near the luminal side increases access of luminal protons to the S_TMD_ site. Protonation of S_TMD_, likely through the two histidine residues, disrupts Zn^2+^ coordination and facilitates Zn^2+^ release from S_TMD_ to the lumen. For better viewing, the other two aspartate residues of the S_TMD_ site (D104/D229) were not displayed.

### Other features of the xtZnT8-Zn^2+^ structure compared to hZnT8-Zn^2+^

Meanwhile, structural comparison between xtZnT8-Zn^2+^ and hZnT8-Zn^2+^ reveals two apparent differences. First, no Zn^2+^ density was observed in the S_interface_ site in xtZnT8-Zn^2+^ (His131/His350) even in the presence of 1 mM Zn^2+^ (Fig. 9A), while the S_interface_ Zn^2+^ has been clearly observed between His137 and His345 in hZnT8-Zn^2+^ (Xue et al., 2020). This difference could be due to different experimental procedures or maybe reflect a less conservation of S_interface_ among ZnT8 species (Fig. 2). Second, the region between the HCH motif and TM1 (HCH-TM1-linker) assumed an α helical structure as a continuation of the TM1 helix in xtZnT8-Zn^2+^. In hZnT8-Zn^2+^, HCH-TMl-linker was not resolved (Fig. 9B), suggesting that this region may adopt a flexible/unordered conformation. It is noteworthy that the distance between the HCH motif and TM1 varies substantially (> 10 Å) between different hZnT8 conformations (Xue et al., 2020), mandating that HCH-TM1-linker is capable of changing conformations. Therefore, the different conformations of HCH-TM1-linker could be a mechanism for regulating conformational changes in eukaryotic ZnT8 (Xue et al., 2020). Further studies will be required to address this question.

**Figure 9.**
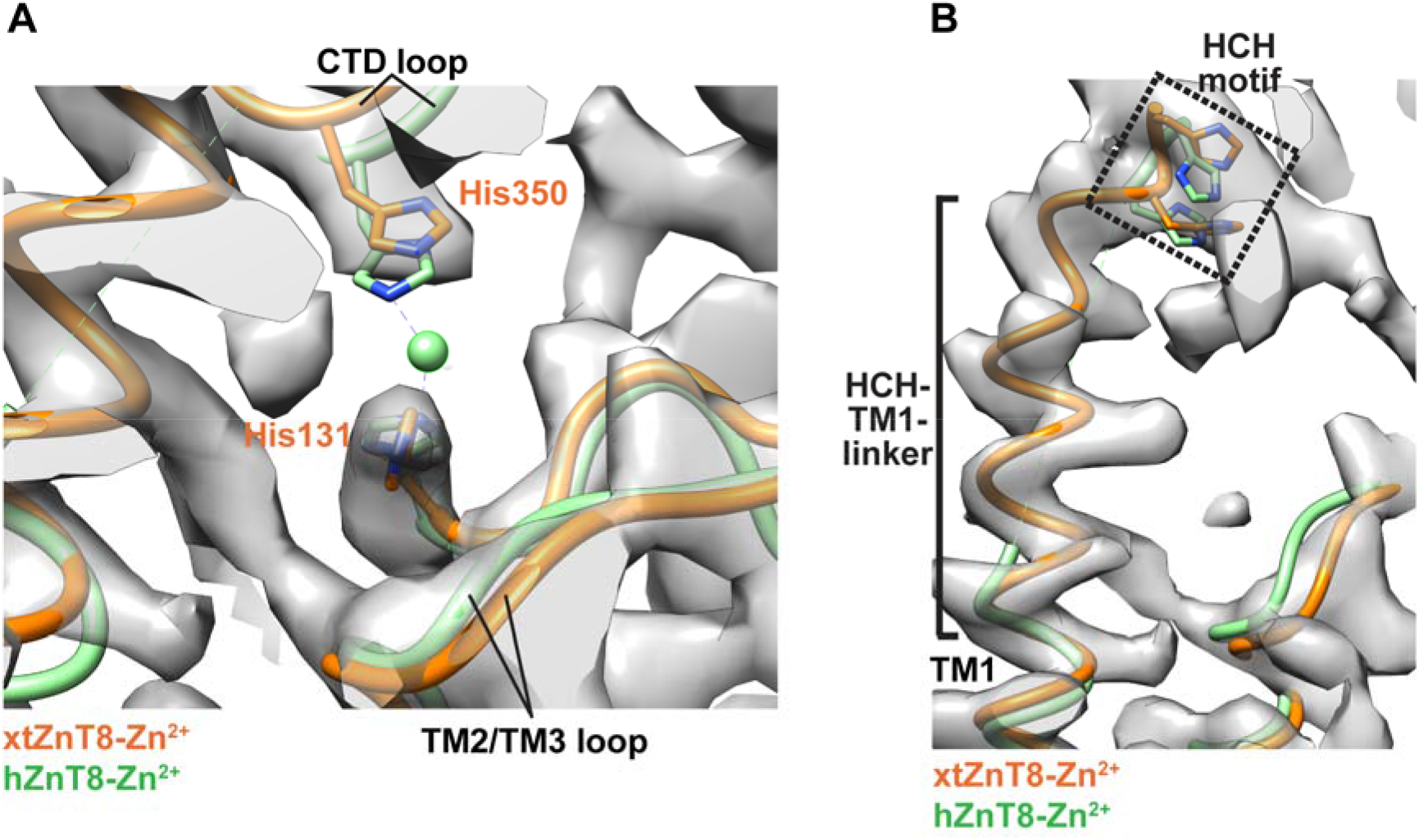
Differences between xtZnT8-Zn^2+^ and hZnT8-Zn^2+^ structures. (A) Comparison of the S_interface_ site. The cryo-EM density map for the S_interface_ site of xtZnT8-Zn^2+^ (orange ribbons) formed by Hisl3l and His35O (orange sticks), corresponding to Hisl37 and His345 (green sticks) in hZnT8-Zn^2+^ (green ribbons). The S_interface_-bound Zn^2+^ in hZnT8-Zn^2+^ is rendered as a green sphere. No Zn^2+^ density was present at the S_interface_ site of xtZnT8-Zn^2+^. (B) The cryo-EM density map of HCH-TM1-linker in xtZnT8-Zn^2+^. The xtZnT8-Zn^2+^ structure is displayed as orange ribbons, and the hZnT8-Zn^2+^ structure (green ribbons) is superimposed onto xtZnT8-Zn^2+^. The HCH motif is boxed and indicated.

## Materials and methods

### Protein expression and purification

The gene encoding the full-length wild-type ZnT8 from *Xenopus tropicalis* (xtZnT8, NCBI accession: NP_001011041.1) was synthesized by Genewiz (Suzhou, China) in a modified pPICZ plasmid (Thermo Fisher Scientific) containing a carboxy-terminal tag of Tobacco etch virus (TEV) protease recognition site and decahistidine. The bacterial cytochrome b562RIL (BRIL) (Chu et al., 2002) was fused to the carboxy-end of the xtZnT8 protein before the TEV site. Mutations were introduced by site-directed mutagenesis using QuikChange II system (Agilent, Santa Clara, CA) according to manufacturer’s recommendation, and all mutations were verified by sequencing. The xtZnT8-BRIL fusion protein (referred to as xtZnT8 in this study) was overexpressed in yeast *(Pichia pastoris* strain GS115) cells by adding 1% (v/v) methanol and 2.5% (v/v) dimethyl sulfoxide (DMSO) at OD_600nm_ of ~1 and shaking at 20 °C for 48 h. Cell pellets were resuspended in lysis solution (LS) containing 20 mM Tris-HCl pH 7.5, 150 mM NaCl, 10% (v/v) glycerol, 1 mM phenylmethanesulfonyl fluoride (PMSF) and 2 mM β-mercaptoethanol, and were lysed by an ATS AH-1500 high-pressure homogenizer (Shanghai, China) at 1,300 MPa. Protein was extracted by addition of 1% (w/v) n-dodecyl-β-D-maltopyranoside (DDM, Anatrace) and 0.1% (w/v) cholesteryl hemisuccinate (CHS, Anatrace) at 4 °C for 2 h and the extraction mixture was centrifuged at 200,000 x g for 20 min at 4 °C. The supernatant was then loaded onto a cobalt metal affinity column, washed with 20 bed-volume of LS containing 1 mM DDM, 0.01% (w/v) CHS and 40 mM imidazole pH 8.0, and eluted with LS supplemented with 1 mM DDM, 0.01% (w/v) CHS and 250 mM imidazole pH 8.0.

### Cryo-EM sample preparation and image acquisition

Affinity-purified xtZnT8 was concentrated to 6-8 mg/ml and loaded onto a Superdex 200 Increase 10/300 GL column (GE Healthcare), and was further purified by size-exclusion chromatography (SEC). For xtZnT8-Zn^2+^, the SEC buffer contained 20 mM Tris-HCl pH 7.5,150 mM NaCl, 5 mM β-mercaptoethanol, 0.5 mM DDM and 1 mM ZnSO_4_. For xtZnT8-H^+^, SEC-purified xtZnT8-Zn^2+^ underwent an additional SEC purification in 20 mM MES-Na pH 5.5, 150 mM NaCl, 5 mM β-mercaptoethanol, 0.5 mM DDM and 1 mM ZnSO_4_. Freshly SEC-purified xtZnT8 was then concentrated to 8 mg/ml as approximated by ultraviolet absorbance, and 3 μl of the protein solution was applied to glow-discharged (45 s) Quantifoil R2/1 300-mesh gold grids (Quantifoil Micro Tools GmbH, Germany). The grids were blotted with standard Vitrobot filter paper for 2.5 s at 4 °C under 100% humidity and plunged into liquid ethane using a Vitrobot Mark IV (Thermo Fisher Scientific). The frozen grids were loaded into a Titan Krios electron microscope (Thermo Fisher Scientific) operated at 300 kV with the condenser lens aperture at 50 μm. Microscope magnification was at 130,000× (1.1 Å per pixel) for xtZnT8-Zn^2+^and 165,000× (0.85 Å per pixel) for xtZnT8-H^+^. Movie stacks were collected automatically using the EPU software (Version 2.9.0.1519REL, Thermo Fisher Scientific) on a K2 Summit direct electron camera (Gatan) equipped with a Quantum GIF energy filter with an energy slit of 20 eV in counting mode. Data were collected at 5 raw frames per second for 6 s, yielding 30 frames per stack and a total exposure dose of 50.1 e^-^/Å^2^ for xtZnT8-Zn^2+^ and 63.9 e^-^/Å^2^ for xtZnT8-H^+^. A total of 2,961 movie stacks for xtZnT8-Zn^2+^ and 2,258 movie stacks for xtZnT8-H^+^ were collected at a defocus range of −1.0 to −1.7 μm.

### Cryo-EM data processing and 3D reconstruction

The movie stacks of xtZnT8-Zn^2+^ were motion-corrected and dose-weighted using the MotionCor2 program (Zheng et al., 2017). CTF correction was performed using the Gctf program (Zhang, 2016). The rest of the data processing was carried out in RELION 3.1 (Zivanov et al., 2018) except the last step. About 20 micrographs from the xtZnT8-Zn^2+^ dataset were used for manual picking of ~1000 particles, which were used for an initial 2D classification. Class averages representing projections of the xtZnT8 dimer in different orientations were selected as templates for automated particle picking from the full dataset. A total of 928,822 particles were picked from 2,943 micrographs. Particles extracted from the dataset were binned by a factor of 3 and subjected to reference-free 2D classification, and particles in good 2D classes were selected (518,886 good particles in total). Using a subset of the selected good particles a coarse initial 3D model was generated first, followed by a one-class (K=1) global 3D classification with all good particles to generate a fine initial 3D model. Using the fine initial 3D model, all good particles were subjected to a four-class (K=4) local 3D classification with an angular search step of 3.75°. The local 3D classification step was performed in six parallel runs, and particles of the best 3D class from each run were combined and duplicate particles were removed, resulting in 380,810 particles. The re-extracted unbinned particles were then subjected to a final six-class multi-reference 3D classification using the best 3D classes from the local 3D classification step as references. The resulting best 3D class containing 233,408 particles was subjected to per-particle defocus refinement, beam-tilt refinement and Bayesian polishing, and was subsequently imported into CryoSPARC 3.2 (Punjani et al., 2017). After Nonuniform refinement (Punjani et al., 2020) in CryoSPARC, the final reconstruction was solved at an overall resolution of 3.9⍰Å. For the xtZnT8-H^+^ dataset, a total of 684,578 particles were picked from 2,230 micrographs. The same image processing procedure was utilized except that a low-pass-filtered map of xtZnT8-Zn^2+^ was used as the initial model for the xtZnT8-H^+^ particles. Eventually, 101,706 particles were selected from the best 3D class, yielding a 3D reconstruction map at 3.7 Å resolution.

### Model building and refinement

Density maps of xtZnT8-Zn^2+^ and xtZnT8-H^+^ were of sufficient quality for de novo model building in Coot (Emsley and Cowtan, 2004), facilitated by a published structure of hZnT8 (PDB: 6XPE). The xtZnT8 models were manually adjusted and rebuilt in Coot and refined using the phenix.real_space_refine module (Afonine et al., 2018) in PHENIX (Adams et al., 2010). Model geometries were assessed by Molprobity (Davis et al., 2007) and summarized in Table 1. Structural figures were generated using PyMOL (Schr⍰dinger, LLC) and UCSF Chimera (Pettersen et al., 2004). Accessibility analysis was performed using the volume-filling program HOLLOW (Ho and Gruswitz, 2008) with default settings.

**Table 1.** Cryo-EM data collection and model statistics.

|  | xtZnT8-Zn<sup>2+</sup><br>(EMD-33619)<br>(PDB: 7Y5G) | xtZnT8-H<sup>+</sup><br>(EMD-33620)<br>(PDB: 7Y5H) |
| --- | --- | --- |
| **Data collection and processing** |  |  |
| Magnification | 130,000 | 165,000 |
| Voltage (kV) | 300 | 300 |
| Electron exposure (e<sup>-</sup>/Å<sup>2</sup>) | 50.1 | 63.9 |
| Defocus range (μm) | -1.0~-1.7 | -1.1~-1.7 |
| Pixel size (Å) | 1.1 | 0.85 |
| Symmetry imposed | C2 | C2 |
| Initial particle images (no.) | 928,822 | 684,578 |
| Final particle images (no.) | 233,408 | 101,706 |
| Map resolution (Å) | 3.85 | 3.72 |
| FSC threshold | 0.143 | 0.143 |
| **Refinement** |  |  |
| Model composition |  |  |
| Non-hydrogen atoms | 4736 | 4704 |
| Protein residues | 604 | 600 |
| Ligands | 6 | 4 |
| R.m.s. deviations |  |  |
| Bond lengths (Å) | 0.002 | 0.002 |
| Bond angles (°) | 0.433 | 0.391 |
| Validation |  |  |
| MolProbity score | 1.01 | 1.16 |
| Clashscore | 2.29 | 3.76 |
| Rotamer outliers (%) | 0.00 | 0.00 |
| Ramachandran plot |  |  |
| Favored (%) | 99.32 | 99.49 |
| Allowed (%) | 0.68 | 0.51 |
| Disallowed (%) | 0.00 | 0.00 |
| *B*-factors (Å<sup>2</sup>) |  |  |
| Protein | 28.11 | 41.22 |
| Ligand | 48.60 | 65.00 |

### Microscale thermophoresis (MST)

MST analysis was performed using Monolith NT.115 (NanoTemper Technologies, Germany) by staining His-tagged xtZnT8 with the RED-Tris-NTA 2nd Generation dye (NanoTemper Technologies, Germany). For Zn^2+^ titration, affinity-purified xtZnT8 was further purified by SEC in a buffer containing 150 mM NaCl, 1 mM DDM, and either 20 mM HEPES-Na pH 7.5 or 20 mM MES-Na pH 5.5 as designed. Peak fractions were pooled and diluted to 200 nM using the SEC buffer, and mixed with 100 nM RED-Tris-NTA 2nd Generation dye at a 1:1 ratio and incubated for 30 min at room temperature. Then the sample was centrifuged at 15,000 x g for 10 min at 4 °C to keep the supernatant containing the labeled protein. Freshly-prepared serial-diluted ZnSO_4_ solutions were used as the titrant and were prepared according to the MST manual with 1 mM as the highest concentration used. The labeled protein was mixed with serial-diluted Zn^2+^ and incubated for 30 min at room temperature. Then the protein sample series were loaded into capillaries and MST measurements were performed according to the Monolith manual. The equilibrium dissociation constant *(K_d_)* was determined using the MO.Affinity Analysis software (NanoTemper Technologies, Germany) with the *K_d_* fit function, and a noise-like trendless pattern for all data points in a measurement that fails the *K_d_* fit function was deemed a no-binding pattern. All MST measurements were performed in three independently repeated experiments, and *K_d_* was expressed as mean ± standard deviation (SD) in the text.

## Abbreviations

Cryo-EM: cryo-electron microscopy
ZnT8: zinc transporter 8
SLC: solute carrier
TMD: transmembrane domain
CTD: carboxy-terminal domain
RMSD: root mean square deviation

## Data Availability

The cryo-EM density maps for xtZnT8-Zn^2+^ and xtZnT8-H^+^ have been deposited in the Electron Microscopy Data Bank under the accession IDs EMD-33619 and EMD-33620, respectively. The atomic coordinates of the xtZnT8-Zn^2+^ and xtZnT8-H^+^ structures have been deposited in the Protein Data Bank under the accession codes 7Y5G and 7Y5H, respectively. Two previously published hZnT8 structures used in this study are available in the Protein Data Bank under accession codes 6XPD and 6XPE.

## Acknowledgements

Cryo-EM data were collected at SKLB West China Cryo-EM Center and were processed at the Duyu High Performance Computing Center of Sichuan University.

## Funding and additional information

This work was supported in part by the National Natural Science Foundation of China (NSFC) grant 31770783 to Xiaoming Zhou and 32070049 to Zhaoming Su, the 1.3.5 Project for Disciplines of Excellence grant by West China Hospital of Sichuan University to Xiaoming Zhou and Zhaoming Su, and Ministry of Science and Technology of China grant 2021YFA1301900 to Zhaoming Su.

## References

Adams, P.D., Afonine, P.V., Bunkoczi, G., Chen, V.B., Davis, I.W., Echols, N., Headd, J.J., Hung, L.W., Kapral, G.J., Grosse-Kunstleve, R.W., McCoy, A.J., Moriarty, N.W., Oeffner, R., Read, R.J., Richardson, D.C., Richardson, J.S., Terwilliger, T.C., Zwart, P.H., 2010. PHENIX: a comprehensive Python-based system for macromolecular structure solution. Acta Crystallogr D Biol Crystallogr 66, 213–221.

Afonine, P.V., Poon, B.K., Read, R.J., Sobolev, O.V., Terwilliger, T.C., Urzhumtsev, A., Adams, P.D., 2018. Real-space refinement in PHENIX for cryo-EM and crystallography. Acta Crystallogr D Struct Biol 74, 531–544.

Bin, B.H., Seo, J., Kim, S.T., 2018. Function, Structure, and Transport Aspects of ZIP and ZnT Zinc Transporters in Immune Cells. J Immunol Res 2018, 9365747.

Chu, R., Takei, J., Knowlton, J.R., Andrykovitch, M., Pei, W., Kajava, A.V., Steinbach, P.J., Ji, X., Bai, Y., 2002. Redesign of a four-helix bundle protein by phage display coupled with proteolysis and structural characterization by NMR and X-ray crystallography. J Mol Biol 323, 253–262.

Combet, C., Blanchet, C., Geourjon, C., Deleage, G., 2000. NPS@: network protein sequence analysis. Trends Biochem Sci 25, 147–150.

Cotrim, C.A., Jarrott, R.J., Martin, J.L., Drew, D., 2019. A structural overview of the zinc transporters in the cation diffusion facilitator family. Acta Crystallogr D Struct Biol 75, 357–367.

Coudray, N., Valvo, S., Hu, M., Lasala, R., Kim, C., Vink, M., Zhou, M., Provasi, D., Filizola, M., Tao, J., Fang, J., Penczek, P.A., Ubarretxena-Belandia, I., Stokes, D.L., 2013. Inward-facing conformation of the zinc transporter YiiP revealed by cryoelectron microscopy. Proc Natl Acad Sci U S A 110, 2140–2145.

Davidson, H.W., Wenzlau, J.M., O’Brien, R.M., 2014. Zinc transporter 8 (ZnT8) and beta cell function. Trends Endocrinol Metab 25, 415–424.

Davis, I.W., Leaver-Fay, A., Chen, V.B., Block, J.N., Kapral, G.J., Wang, X., Murray, L.W., Arendall, W.B., 3rd, Snoeyink, J., Richardson, J.S., Richardson, D.C., 2007. MolProbity: all-atom contacts and structure validation for proteins and nucleic acids. Nucleic Acids Res 35, W375–383.

Eide, D.J., 2006. Zinc transporters and the cellular trafficking of zinc. Biochim Biophys Acta 1763, 711–722.

Emsley, P., Cowtan, K., 2004. Coot: model-building tools for molecular graphics. Acta Crystallogr D Biol Crystallogr 60, 2126–2132.

Flannick, J., Thorleifsson, G., Beer, N.L., Jacobs, S.B., Grarup, N., Burtt, N.P., Mahajan, A., Fuchsberger, C., Atzmon, G., Benediktsson, R., Blangero, J., Bowden, D.W., Brandslund, I., Brosnan, J., Burslem, F., Chambers, J., Cho, Y.S., Christensen, C., Douglas, D.A., Duggirala, R., Dymek, Z., Farjoun, Y., Fennell, T., Fontanillas, P., Forsen, T., Gabriel, S., Glaser, B., Gudbjartsson, D.F., Hanis, C., Hansen, T., Hreidarsson, A.B., Hveem, K., Ingelsson, E., Isomaa, B., Johansson, S., Jorgensen, T., Jorgensen, M.E., Kathiresan, S., Kong, A., Kooner, J., Kravic, J., Laakso, M., Lee, J.Y., Lind, L., Lindgren, C.M., Linneberg, A., Masson, G., Meitinger, T., Mohlke, K.L., Molven, A., Morris, A.P., Potluri, S., Rauramaa, R., Ribel-Madsen, R., Richard, A.M., Rolph, T., Salomaa, V., Segre, A.V., Skarstrand, H., Steinthorsdottir, V., Stringham, H.M., Sulem, P., Tai, E.S., Teo, Y.Y., Teslovich, T., Thorsteinsdottir, U., Trimmer, J.K., Tuomi, T., Tuomilehto, J., Vaziri-Sani, F., Voight, B.F., Wilson, J.G., Boehnke, M., McCarthy, M.I., Njolstad, P.R., Pedersen, O., Go, T.D.C., Consortium, T.D.G., Groop, L., Cox, D.R., Stefansson, K., Altshuler, D., 2014. Loss-of-function mutations in SLC30A8 protect against type 2 diabetes. Nat Genet 46, 357–363.

Gupta, S., Chai, J., Cheng, J., D’Mello, R., Chance, M.R., Fu, D., 2014. Visualizing the kinetic power stroke that drives proton-coupled zinc(ll) transport. Nature 512, 101–104.

Ho, B.K., Gruswitz, F., 2008. HOLLOW: generating accurate representations of channel and interior surfaces in molecular structures. BMC Struct Biol 8, 49.

Hoch, E., Lin, W., Chai, J., Hershfinkel, M., Fu, D., Sekler, I., 2012. Histidine pairing at the metal transport site of mammalian ZnT transporters controls Zn2+ over Cd2+ selectivity. Proc Natl Acad Sci U S A 109, 7202–7207.

Hutton, J.C., 1982. The internal pH and membrane potential of the insulin-secretory granule. Biochem J 204, 171–178.

Jerabek-Willemsen, M., Wienken, C.J., Braun, D., Baaske, P., Duhr, S., 2011. Molecular interaction studies using microscale thermophoresis. Assay Drug Dev Technol 9, 342–353.

Kambe, T., Tsuji, T., Hashimoto, A., Itsumura, N., 2015. The Physiological, Biochemical, and Molecular Roles of Zinc Transporters in Zinc Homeostasis and Metabolism. Physiol Rev 95, 749–784.

Kawasaki, E., 2012. ZnT8 and type 1 diabetes. Endocr J 59, 531–537.

Llopis, J., McCaffery, J.M., Miyawaki, A., Farquhar, M.G., Tsien, R.Y., 1998. Measurement of cytosolic, mitochondrial, and Golgi pH in single living cells with green fluorescent proteins. Proc Natl Acad Sci U S A 95, 6803–6808.

Lopez-Redondo, M., Fan, S., Koide, A., Koide, S., Beckstein, O., Stokes, D.L., 2021. Zinc binding alters the conformational dynamics and drives the transport cycle of the cation diffusion facilitator YiiP. J Gen Physiol 153.

Lopez-Redondo, M.L., Coudray, N., Zhang, Z., Alexopoulos, J., Stokes, D.L., 2018. Structural basis for the alternating access mechanism of the cation diffusion facilitator YiiP. Proc Natl Acad Sci U S A 115, 3042–3047.

Lu, M., Fu, D., 2007. Structure of the zinc transporter YiiP. Science 317, 1746–1748.

Lu, M., Chai, J., Fu, D., 2009. Structural basis for autoregulation of the zinc transporter YiiP. Nat Struct Mol Biol 16, 1063–1067.

Maret, W., 2013. Zinc biochemistry: from a single zinc enzyme to a key element of life. Adv Nutr 4, 82–91.

Ohana, E., Hoch, E., Keasar, C., Kambe, T., Yifrach, O., Hershfinkel, M., Sekler, I., 2009. Identification of the Zn2+ binding site and mode of operation of a mammalian Zn2+ transporter. J Biol Chem 284, 17677–17686.

Pettersen, E.F., Goddard, T.D., Huang, C.C., Couch, G.S., Greenblatt, D.M., Meng, E.C., Ferrin, T.E., 2004. UCSF Chimera--a visualization system for exploratory research and analysis. J Comput Chem 25, 1605–1612.

Punjani, A., Zhang, H., Fleet, D.J., 2020. Non-uniform refinement: adaptive regularization improves single-particle cryo-EM reconstruction. Nat Methods 17, 1214–1221.

Punjani, A., Rubinstein, J.L., Fleet, D.J., Brubaker, M.A., 2017. cryoSPARC: algorithms for rapid unsupervised cryo-EM structure determination. Nat Methods 14, 290–296.

Saxena, R., Voight, B.F., Lyssenko, V., Burtt, N.P., de Bakker, P.I., Chen, H., Roix, J.J., Kathiresan, S., Hirschhorn, J.N., Daly, M.J., Hughes, T.E., Groop, L., Altshuler, D., Almgren, P., Florez, J.C., Meyer, J., Ardlie, K., Bengtsson Bostrom, K., Isomaa, B., Lettre, G., Lindblad, U., Lyon, H.N., Melander, O., Newton-Cheh, C., Nilsson, P., Orho-Melander, M., Rastam, L., Speliotes, E.K., Taskinen, M.R., Tuomi, T., Guiducci, C., Berglund, A., Carlson, J., Gianniny, L., Hackett, R., Hall, L., Holmkvist, J., Laurila, E., Sjogren, M., Sterner, M., Surti, A., Svensson, M., Svensson, M., Tewhey, R., Blumenstiel, B., Parkin, M., Defelice, M., Barry, R., Brodeur, W., Camarata, J., Chia, N., Fava, M., Gibbons, J., Handsaker, B., Healy, C., Nguyen, K., Gates, C., Sougnez, C., Gage, D., Nizzari, M., Gabriel, S.B., Chirn, G.W., Ma, Q., Parikh, H., Richardson, D., Rieke, D., Purcell, S., 2007. Genome-wide association analysis identifies loci for type 2 diabetes and triglyceride levels. Science 316, 1331–1336.

Scott, L.J., Mohlke, K.L., Bonnycastle, L.L., Willer, C.J., Li, Y., Duren, W.L., Erdos, M.R., Stringham, H.M., Chines, P.S., Jackson, A.U., Prokunina-Olsson, L., Ding, C.J., Swift, A.J., Narisu, N., Hu, T., Pruim, R., Xiao, R., Li, X.Y., Conneely, K.N., Riebow, N.L., Sprau, A.G., Tong, M., White, P.P., Hetrick, K.N., Barnhart, M.W., Bark, C.W., Goldstein, J.L., Watkins, L., Xiang, F., Saramies, J., Buchanan, T.A., Watanabe, R.M., Valle, T.T., Kinnunen, L., Abecasis, G.R., Pugh, E.W., Doheny, K.F., Bergman, R.N., Tuomilehto, J., Collins, F.S., Boehnke, M., 2007. A genome-wide association study of type 2 diabetes in Finns detects multiple susceptibility variants. Science 316, 1341–1345.

Shusterman, E., Beharier, O., Shiri, L., Zarivach, R., Etzion, Y., Campbell, C.R., Lee, I.H., Okabayashi, K., Dinudom, A., Cook, D.I., Katz, A., Moran, A., 2014. ZnT-1 extrudes zinc from mammalian cells functioning as a Zn(2+)/H(+) exchanger. Metallomics 6, 1656–1663.

Sladek, R., Rocheleau, G., Rung, J., Dina, C., Shen, L., Serre, D., Boutin, P., Vincent, D., Belisle, A., Hadjadj, S., Balkau, B., Heude, B., Charpentier, G., Hudson, T.J., Montpetit, A., Pshezhetsky, A.V., Prentki, M., Posner, B.I., Balding, D.J., Meyre, D., Polychronakos, C., Froguel, P., 2007. A genome-wide association study identifies novel risk loci for type 2 diabetes. Nature 445, 881–885.

Stiernet, P., Nenquin, M., Moulin, P., Jonas, J.C., Henquin, J.C., 2007. Glucose-induced cytosolic pH changes in beta-cells and insulin secretion are not causally related: studies in islets lacking the Na+/H+ exchangeR NHE1. J Biol Chem 282, 24538–24546.

Thompson, J.D., Higgins, D.G., Gibson, T.J., 1994. CLUSTAL W: improving the sensitivity of progressive multiple sequence alignment through sequence weighting, position-specific gap penalties and weight matrix choice. Nucleic Acids Res 22, 4673–4680.

Vallee, B.L., Falchuk, K.H., 1993. The biochemical basis of zinc physiology. Physiol Rev 73, 79–118.

Wenzlau, J.M., Juhl, K., Yu, L., Moua, O., Sarkar, S.A., Gottlieb, P., Rewers, M., Eisenbarth, G.S., Jensen, J., Davidson, H.W., Hutton, J.C., 2007. The cation efflux transporter ZnT8 (Slc30A8) is a major autoantigen in human type 1 diabetes. Proc Natl Acad Sci U S A 104, 17040–17045.

Xue, J., Xie, T., Zeng, W., Jiang, Y., Bai, X.C., 2020. Cryo-EM structures of human ZnT8 in both outward-and inward-facing conformations. Elife 9.

Zeggini, E., Weedon, M.N., Lindgren, C.M., Frayling, T.M., Elliott, K.S., Lango, H., Timpson, N.J., Perry, J.R., Rayner, N.W., Freathy, R.M., Barrett, J.C., Shields, B., Morris, A.P., Ellard, S., Groves, C.J., Harries, L.W., Marchini, J.L., Owen, K.R., Knight, B., Cardon, L.R., Walker, M., Hitman, G.A., Morris, A.D., Doney, A.S., Wellcome Trust Case Control, C., McCarthy, M.I., Hattersley, A.T., 2007. Replication of genome-wide association signals in UK samples reveals risk loci for type 2 diabetes. Science 316, 1336–1341.

Zhang, K., 2016. Gctf: Real-time CTF determination and correction. J Struct Biol 193, 1–12.

Zheng, S.Q., Palovcak, E., Armache, J.P., Verba, K.A., Cheng, Y., Agard, D.A., 2017. MotionCor2: anisotropic correction of beam-induced motion for improved cryo-electron microscopy. Nat Methods 14, 331–332.

Zivanov, J., Nakane, T., Forsberg, B.O., Kimanius, D., Hagen, W.J., Lindahl, E., Scheres, S.H., 2018. New tools for automated high-resolution cryo-EM structure determination in RELION-3. Elife 7.

